# Structural and Evolutionary Analysis Indicate that the SARS-CoV-2 Mpro is an Inconvenient Target for Small-Molecule Inhibitors Design

**DOI:** 10.1101/2020.02.27.968008

**Authors:** Maria Bzówka, Karolina Mitusińska, Agata Raczyńska, Aleksandra Samol, Jack A. Tuszyński, Artur Góra

## Abstract

The novel coronavirus whose outbreak took place in December 2019 continues to spread at a rapid rate worldwide. In the absence of an effective vaccine, inhibitor repurposing or de novo drug design may offer a longer-term strategy to combat this and future infections due to similar viruses. Here, we report on detailed classical and mix-solvent molecular dynamics simulations of the main protease (Mpro) enriched by evolutionary and stability analysis of the protein. The results were compared with those for a highly similar SARS Mpro protein. In spite of a high level of sequence similarity, the active sites in both proteins show major differences in both shape and size indicating that repurposing SARS drugs for COVID-19 may be futile. Furthermore, analysis of the binding site’s conformational changes during the simulation time indicates its flexibility and plasticity, which dashes hopes for rapid and reliable drug design. Conversely, structural stability of the protein with respect to flexible loop mutations indicates that the virus’ mutability will pose a further challenge to the rational design of small-molecule inhibitors. However, few residues contribute significantly to the protein stability and thus can be considered as key anchoring residues for Mpro inhibitor design.

## 1. Introduction

In early December 2019, the first atypical pneumonia outbreak associated with the novel coronavirus of zoonotic origin (SARS-CoV-2) appeared in Wuhan City, Hubei Province, China [1,2]. In general, coronaviruses (CoVs) are classified into four major genera: Alphacoronavirus, Betacoronavirus (which primarily infect mammals), Gammacoronavirus, and Deltacoronavirus (which primarily infect birds) [3–5]. In humans, coronaviruses usually cause mild to moderate upper-respiratory tract illnesses, e.g., the common cold, however, the rarer forms of CoVs can be lethal. By the end of 2019, six kinds of human CoV have been identified: HCoV-NL63, HCoV-229E, belonging to Alphacoronavirus genera, HCoV-OC43, HCoV-HKU1, severe acute respiratory syndrome SARS-CoV, and Middle East respiratory syndrome MERS-CoV, belonging to Betacoronavirus genera [4]. Of the aforementioned CoVs, the last two are the most dangerous and they were associated with the outbreak of two epidemics at the beginning of the 21st century [6]. On January 2020, SARS-CoV-2 was isolated and announced as a new, seventh, type of human coronavirus. It was classified as Betacoronavirus [2]. Based on the phylogenetic analysis of the genomic data of SARS-CoV-2, Zhang et al. indicated that the SARS-CoV-2 is most closely related to two SARS-CoV sequences isolated from bats in 2015 and 2017. This is suggestive that the bat’s CoV and SARS-CoV-2 share a common ancestor, and the new virus can be considered as a SARS-like virus [7].

The genome of coronaviruses typically contains a positive-sense, single-stranded RNA but it differs in size ranging between ~26 and ~32 kb. It also includes a variable number of open reading frames (ORFs) – from 6 to 11. The first ORF is the largest, encoding nearly 70% of the entire genome and 16 non-structural proteins (nsps) [3,8]. Of the nsps, the main protease (Mpro, also known as a chymotrypsin-like cysteine protease, 3CLpro), encoded by nsp5, has been found to play a fundamental role in viral gene expression and replication, thus it is an attractive target for anti-CoV drug design [9]. The remaining ORFs encode accessory and structural proteins, including spike surface glycoprotein (S), small envelope protein (E), matrix protein (M), and nucleocapsid protein (N).

Based on the three sequenced genomes of SARS-CoV-2 (Wuhan/IVDC-HB-01/2019, Wuhan/IVDC-HB-04/2019, and Wuhan/IVDC-HB-05/2019, provided by the National Institute for Viral Disease Control and Prevention, CDC, China), Wu et al., performed a detailed genome annotation. The results were further compared to related coronaviruses – 1,008 human SARS-CoV, 338 bat SARS-like CoV, and 3,131 human MERS-CoV indicating that the three strains of SARS-CoV-2 have almost identical genomes with 14 ORFs, encoding 27 proteins including 15 non-structural proteins (nsp1-10 and nsp12-16), 4 structural proteins (S, E, M, N), and 8 accessory proteins (3a, 3b, p6, 7a, 7b, 8b, 9b, and orf14). The only identified difference in the genome consisting of ~29.8 kb nucleotides consisted of five nucleotides. The genome annotation revealed that SARS-CoV-2 is fairly similar to SARS-CoV at the amino acid level, however, there are some differences in the occurrence of accessory proteins, e.g., the 8a accessory protein, present in SARS-CoV, is absent in SARS-CoV-2 and the lengths of 8b and 3b proteins do not match. The phylogenetic analysis of SARS-CoV-2 showed it to be most closely related to SARS-like bat viruses, but no strain of SARS-like bat virus was found to cover all equivalent proteins of SARS-CoV-2 [10].

As previously mentioned, the main protease is one of the key enzymes in the viral life cycle. Together with other non-structural proteins (papain-like protease, helicase, RNA-dependent RNA polymerase) and the spike glycoprotein structural protein, it is essential for interactions between the virus and host cell receptor during viral entry [11]. Initial analyses of genomic sequences of the four nsps mentioned above indicate that those enzymes are highly conserved sharing more than 90% sequence similarity with the corresponding SARS-CoV enzymes [12].

The first released crystal structure of the Mpro of SARS-CoV-2 (PDB ID: 6lu7) was obtained by Prof. Yang’s group from ShanghaiTech by co-crystallisation with a peptide-like inhibitor N-[(5 methylisoxazol-3-yl)carbonyl]alanyl-L-valyl-N~1-((1R,2Z)-4-(benzyloxy)-4-oxo-1-{[(3R)-2-oxopyrrolidin-3-yl]methyl}but-2-enyl)-L-leucinamide (N3 or PRD_002214) [13]. The same inhibitor was co-crystallised with other human coronaviruses, e.g., HCoV-NL63 (PDB ID: 5gwy), HCoV-KU1 (PDB ID: 3d23), or SARS-CoV (PDB ID: 2amq). This enzyme naturally forms a dimer whose each monomer consists of the N-terminal catalytic region and a C-terminal region [14]. While 12 residues differ between both CoVs, only one, namely S46 in SARS-CoV-2 (A46 in SARS-CoV), is located in the proximity of the entrance to the active site. However, such a small structural change would typically be not expected to substantially affect the binding of small molecules [12]. Such an assumption would routinely involve the generation of a library of derivatives and analogous based on the scaffold of a drug that inhibits the corresponding protein in the SARS-CoV case. As shown in the present paper, regrettably, this strategy is not likely to succeed with SARS-CoV-2 for Mpro as a molecular target.

In this study, we investigate how only 12 different residues, located mostly on the protein’s surface, may affect the behaviour of the active site pocket of the SARS-CoV-2 Mpro structure. To this end, we performed classical molecular dynamics simulations (cMD) of both SARS and SARS-CoV-2 Mpros as well as mixed-solvents molecular dynamics simulations (MixMD) combined with small molecules’ tracking approach to analyse the conformational changes in the binding site. In spite of the structural differences in the active sites of both Mpro proteins, major issues involving plasticity and flexibility of the binding site could result in significant difficulties in inhibitor design for this molecular target. Indeed, an in silico attempt has already been made involving a massive virtual screening for Mpro inhibitors of SARS-CoV-2 using Deep Docking [15]. Other recent attempts focused on virtual screening for putative inhibitors of the same main protease of SARS-CoV-2 based on the clinically approved drugs [16–21], and also based on the compounds from different databases or libraries [22–24]. However, none of such attempts is likely to lead to clinical advances in the fight against SARS-CoV-2 for reasons we elaborate below.

## 2. Results

### 2.1. Crystal structure comparison, and location of the replaced amino acids distal to the active site

The first SARS-CoV-2 main protease’s crystallographic structure was made publicly available through the Protein Data Bank (PDB) [25] as a complex with an N3 inhibitor (PDB ID: 6lu7) [13]. Next, the structure without the inhibitor was also made available (PDB ID: 6y2e) [26]. We refer to those structures as SARS-CoV-2 Mpro^N3^ and SARS-CoV-2 Mpro, respectively. We also used two structures of the SARS-CoV main protease: one, referred to as SARS-CoV Mpro^N3^ (PDB ID: 2amq), co-crystallised with the same N3 inhibitor, and the other without an inhibitor (PDB ID: 1q2w), which we refer to as SARS-CoV Mpro. The SARS-CoV-2 Mpro and SARS-CoV Mpro structures differ by only 12 amino acids located mostly on the proteins’ surface (Figure 1A, Supplementary Table S1). Both enzymes share the same structural composition; they comprise three domains: domains I (residues 1-101) and II (residues 102-184) consist of an antiparallel β-barrel, and the α-helical domain III (residues 201-301) is required for the enzymatic activity [27]. Both enzymes resemble the structure of cysteine proteases, although their active site is lacking the third catalytic residue [28]; their active site comprises a catalytic dyad, namely H41 and C145, and a particularly stable water molecule forms at least three hydrogen bond interactions with surrounding residues, including the catalytic histidine, which corresponds to the position of a third catalytic member (Figure 1B). It should be also noted that one of the differing amino acids in SARS-CoV-2 Mpro, namely S46, is located on a C44-P52 loop, which is flanking the active site cavity.

**Figure 1.**
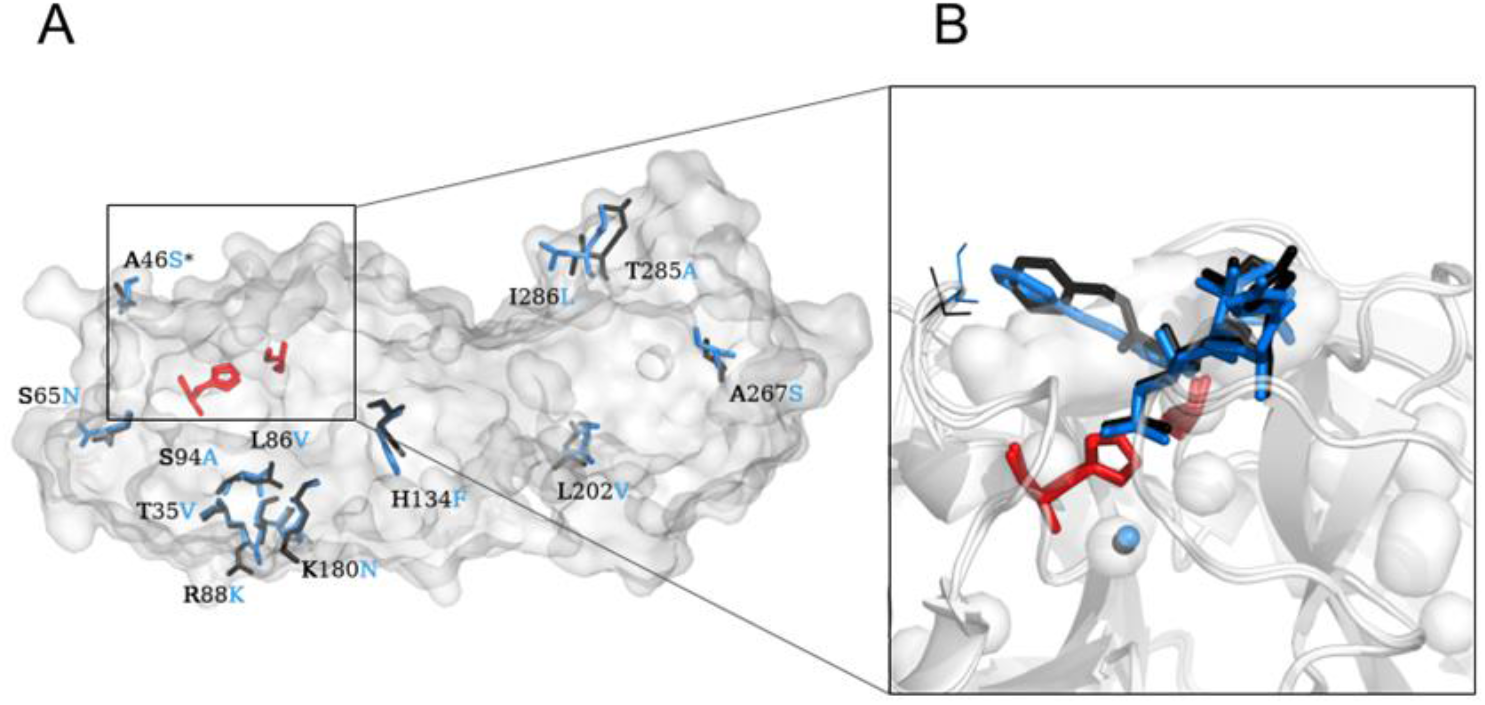
The differences between the SARS-CoV Mpro and SARS-CoV-2 Mpro structures. (A) The overall structure of both SARS-CoV and SARS-CoV-2 Mpros with differing amino acids marked as black (SARS-CoV Mpro) and blue (SARS-CoV-2 Mpro). (B) Close-up of the active site cavity and bound N3 inhibitor into SARS-CoV (black sticks) and SARS-CoV-2 (blue sticks) Mpros. The catalytic water molecule that resembles the position of the third member of the catalytic triad adopted from the cysteine proteases is shown for both SARS-CoV (black sphere) and SARS-CoV-2 (blue sphere) Mpros. The active site residues are shown as red sticks and the proteins’ structures are shown in surface representation. The differing residues in position 46 located near the entrance to the active site are marked with an asterisk (*) on the (A) and as blue and black lines on the (B) panel.

### 2.2. Plasticity of the binding cavities

A total of 2 μs classical molecular dynamics (cMD) simulations of both SARS-CoV-2 and SARS-CoV Mpros with different starting points were run to examine the plasticity of their binding cavities. A combination of the cMD approach with water molecules used as molecular probes is assumed to provide a highly detailed picture of the protein’s interior dynamics [29]. The small molecules tracking approach was used to determine the accessibility of the active site pocket in both SARS-CoV and SARS-CoV-2 Mpros, and a local distribution approach was used to provide information about an overall distribution of solvent in the proteins’ interior. To properly examine the flexibility of both active site cavities, we used the time-window mode implemented in AQUA-DUCT software [30] to analyse the water molecules’ flow through the cavity in a 10 ns time step and combined that with the outer pocket calculations to examine the plasticity and maximal accessible volume (MAV) of the binding cavity.

Surprisingly, despite their high similarity, the binding cavities of SARS-CoV and SARS-CoV-2 Mpros show significantly different MAV. Both proteins reduce their MAV upon inhibitor binding by approximately 20%, but the maximal volume of SARS-CoV is over 50% larger than those of SARS-CoV-2 (Figure 2, Supplementary Figure S1).

**Figure 2.**
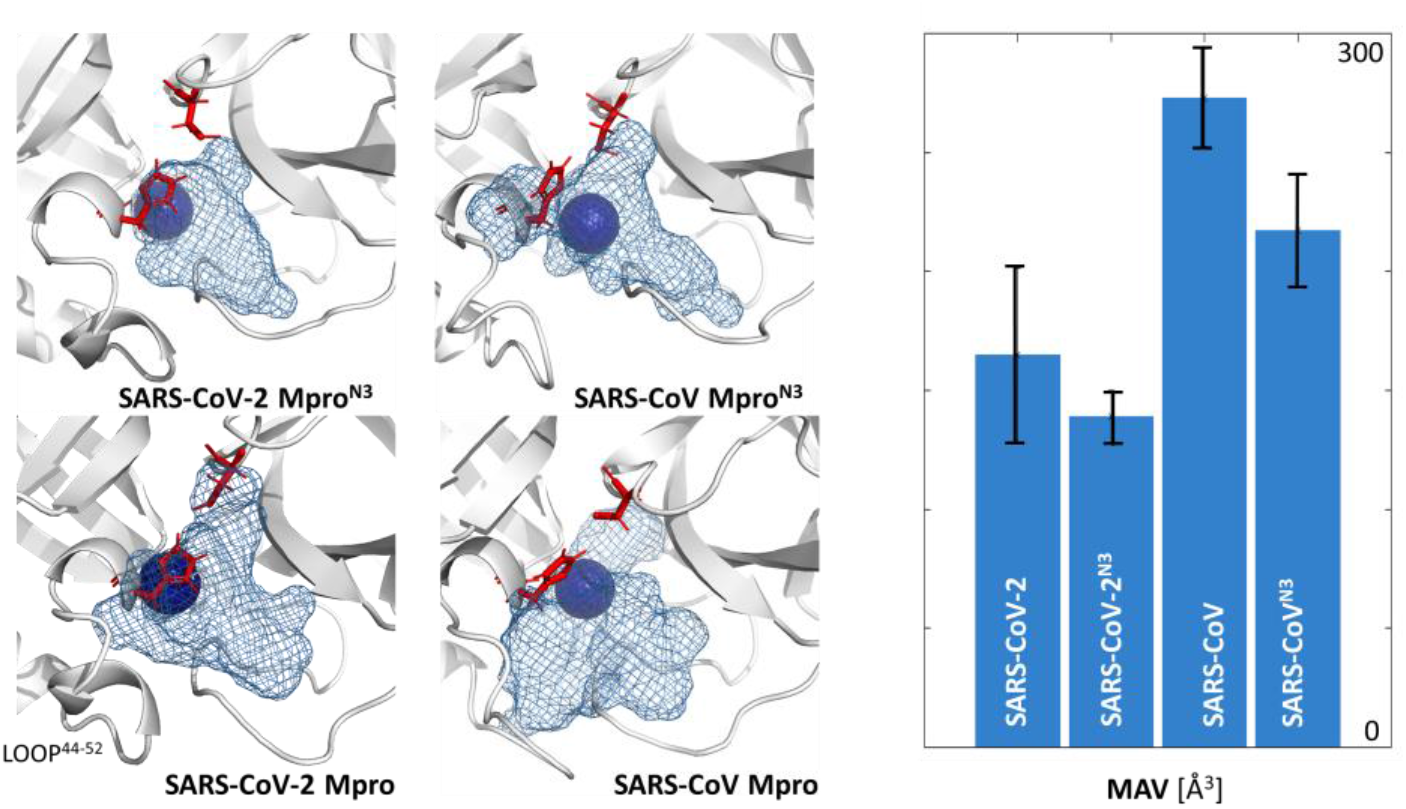
The differences between the maximal accessible volume of the binding cavities calculated during MD simulations of both apo structures of Mpros (SARS-CoV and SARS-CoV-2) and structures with co-crystallised N3 inhibitor (SARS-CoV^N3^ and SARS-CoV-2^N3^) used as different starting points.

### 2.3. Flexibility of the active site entrance

To further examine the plasticity and flexibility of the main proteases binding cavities, we focused on the movements of loops surrounding their entrances and regulating the active sites’ accessibility. We found that one of the analysed loops of the SARS-CoV Mpro, namely C44-P52 loop, is more flexible than the corresponding loops of SARS-CoV-2 Mpro structure, while the adjacent loops are mildly flexible (Figure 3). This could be indirectly assumed from the absence of the C44-P52 loop in the crystallographic structure of SARS-CoV Mpro structure. On the other hand, such flexibility could suggest that the presence of an inhibitor might stabilise the loops surrounding the active site. It is worth adding, that this loop is carrying the unique SARS-CoV-2 Mpro residue S46.

**Figure 3.**
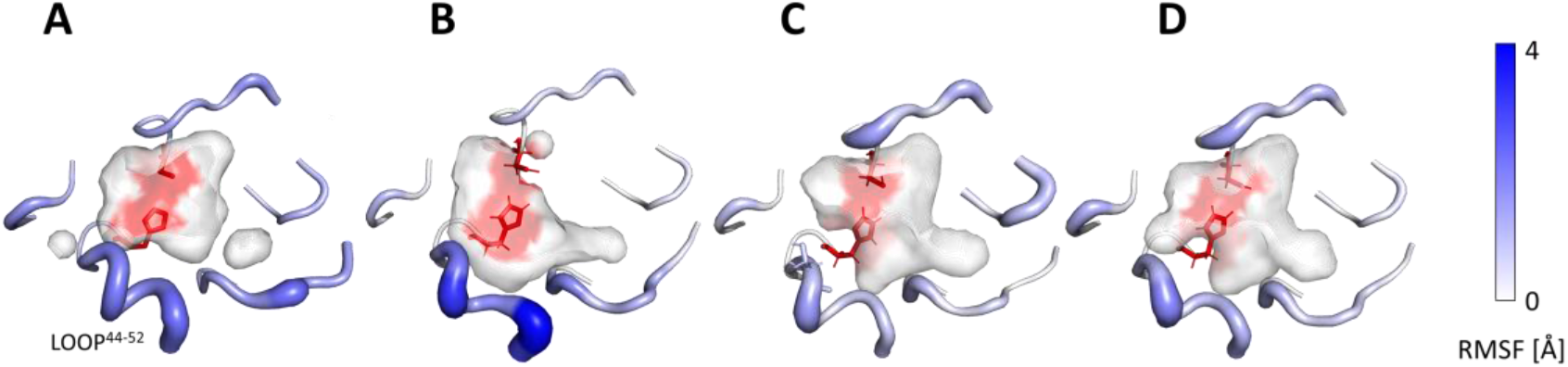
Flexibility of loops surrounding the entrance to the binding cavity of (A) SARS-CoV-2 Mpro, (B) SARS-CoV Mpro, (C) SARS-CoV Mpro^N3^, and (D) SARS-CoV Mpro^N3^. For the picture clarity, only residues creating loops were shown. The active site residues are shown as red sticks and the A46S replacement between SARS-CoV and SARS-CoV-2 main proteases is shown as light blue sticks. The width and colour of the shown residues reflect the level of loop flexibility. The wider and darker residues are more flexible.

### 2.4. Cosolvent hot-spots analysis

The mixed-solvent MD simulations were run with 6 cosolvents: acetonitrile (ACN), benzene (BNZ), dimethylsulfoxide (DMSO), methanol (MEO), phenol (PHN), and urea (URE). Cosolvents were used as specific molecular probes, representing different chemical properties and functional groups that would complement the different regions of the binding site and the protein itself. Using small molecules tracking approach we analysed the flow through the Mpros structures and identified the regions in which those molecules are being trapped and/or caged, located within the protein itself (global hot-spots; Figure 4, Supplementary Figure S2) and inside the binding cavity (local hot-spots; Figure 5, Supplementary Figure S3). The size and location of both types of hot-spots differ and provide complementary information. The global hot-spots identify potential binding/interacting sites in the whole protein structure and additionally provide information about regions attracting particular types of molecules, whereas local hot-spots describe the actual available binding space of a specific cavity.

**Figure 4.**
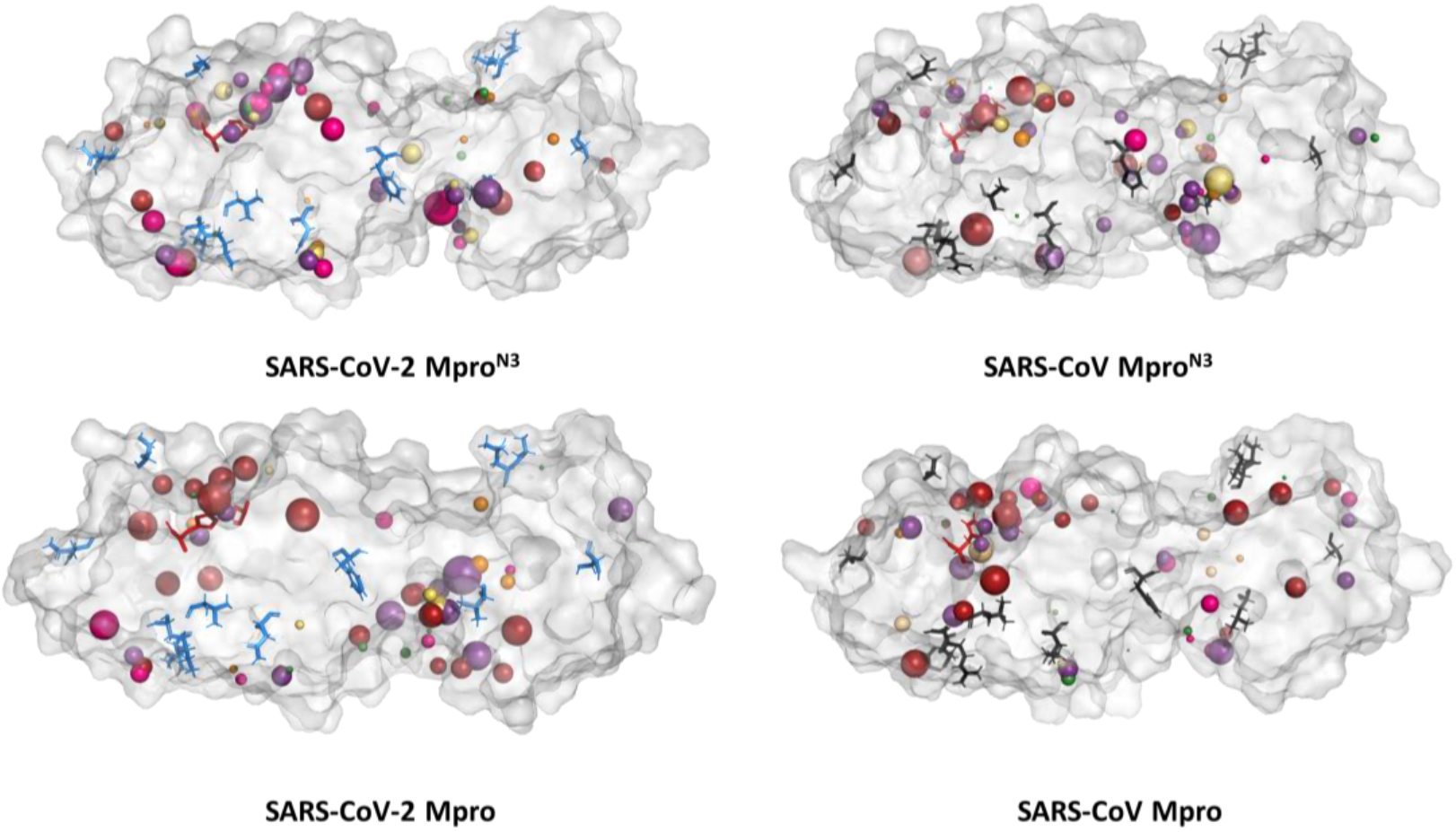
Localisation of the global hot-spots identified in the binding site cavities in SARS-CoV-2 and SARS-CoV main proteases. Hot-spots of individual cosolvents are represented by spheres, and their size reflects the hot-spots density. The colour coding is as follows: purple - urea, green - dimethylsulfoxide, yellow - methanol, orange - acetonitrile, pink - phenol, red - benzene.

The general distribution of the global hot-spots from particular cosolvents is quite similar and verifies specific interactions with the particular regions of the analysed proteins. A notable number of hot-spots are located around the amino acids that vary between the SARS-CoV-2 and SARS-CoV Mpros (Figure 4, Supplementary Figure S2). The largest number and the densest hot-spots are located within the binding cavity and the region essential for Mpros dimerisation [31], between the II and III domains. The binding cavity is particularly occupied by urea, benzene and phenol hot-spots, which is especially interesting, since these solvents exhibit different chemical properties.

A close inspection of the binding site cavity provides further details of cosolvent distribution. The benzene hot-spots for the SARS-CoV-2 Mpro structure are localised deep inside the active site cavity, while SARS-CoV Mpro features mostly benzene hot-spots at the cavity entrance (Figure 5). This is interesting since, in the absence of cosolvent molecules, the water accessible volume for SARS-CoV-2 Mpro was 50% smaller than in the case of SARS-CoV Mpro, underlining huge plasticity of the binding cavity and suggesting large conformational changes induced by interaction with a potential ligand. It is also interesting, that both global and local hot-spots of the SARS-CoV Mpro structure are located in the proximity of the C44-P52 loop, which potentially regulates the access to the active site.

**Figure 5.**
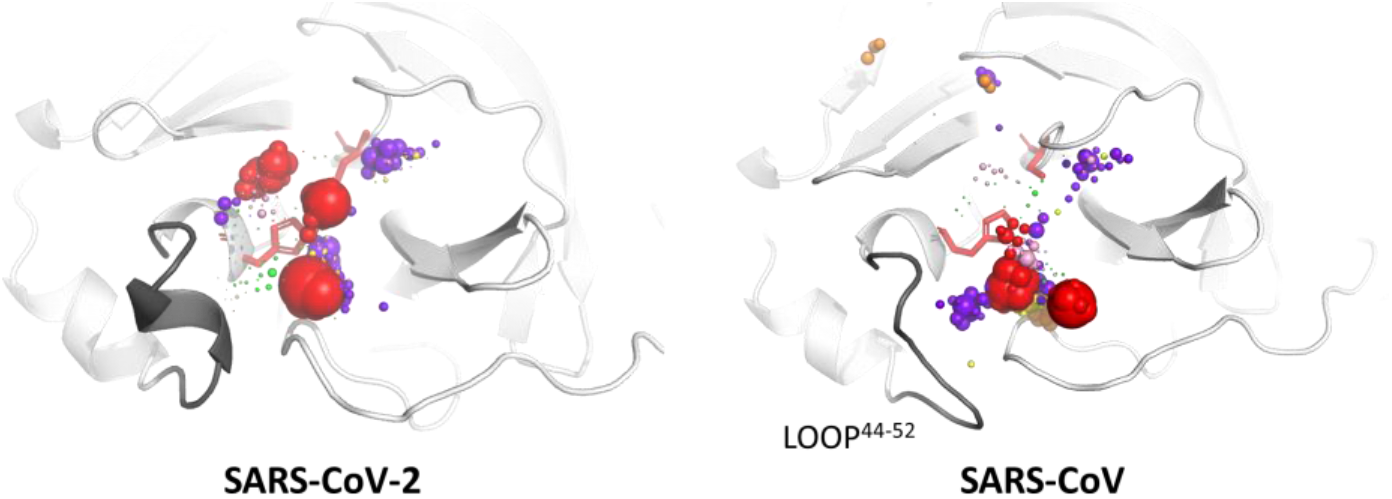
Localisation of the local hot-spots identified in the binding site cavities in SARS-CoV-2 and SARS-CoV main proteases. Hot-spots of individual cosolvents are represented by spheres, and their size reflects the hot-spots density. The colour coding is as follows: purple - urea, green - dimethylsulfoxide, yellow - methanol, orange - acetonitrile, pink - phenol, red - benzene. The active site residues are shown as red sticks, and the proteins’ structures are shown in cartoon representation, loop 44-52 is grey.

### 2.5. Potential mutability of SARS-CoV-2

In general, all the above-mentioned findings indicate potential difficulties in the identification of specific inhibitors toward Mpro proteins. First, the binding site itself is characterised by huge plasticity and probably even distant to active site mutations modify Mpro binding properties. Secondly, the C44-P52 loop regulates access to the active site and can contribute to the discrimination of potential inhibitors. Therefore, additional mutations in the above-mentioned regions, which could appear during further SARS-CoV-2 evolution, can significantly change the affinity between Mpro and its ligands. To verify potential threat of further mutability of the Mpro protein we performed: i) correlated mutation analysis (CMA) on multiple sequence alignment, ii) the analysis of the contribution of already identified differences between the SARS-CoV and SARS-CoV-2 Mpros to protein stability, and iii) prediction of further possible mutations caused by the most probable mutations, the substitution of single nucleotides in the mRNA sequence of Mpro.

Indeed, the analysis performed with Comulator software [32] shows that within viral Mpros evolutionary-correlated residues are dispersed throughout the structure. This indirectly supports our previous findings that distant amino acids mutation can contribute significantly to the binding site plasticity. It is worth adding that among evolutionary-correlated residues we identified also those that differ between SARS-CoV-2 and SARS-CoV Mpros, located on the C44-P52 loop (Figure 6) and the F185-T201 linker loop. The CMA analysis indicates that particular residues in both loops are evolutionary-correlated. The Q189 from the linker loop correlates with residues from the C44-P52 loop, whereas R188, A191, and A194 correlate with selected residues from all domains, but not with the C44-P52 loop. As was shown, the mutation of amino acids distant from active site residues, that are evolutionary correlated, is most likely to modify the active site accessibility [33].

**Figure 6.**
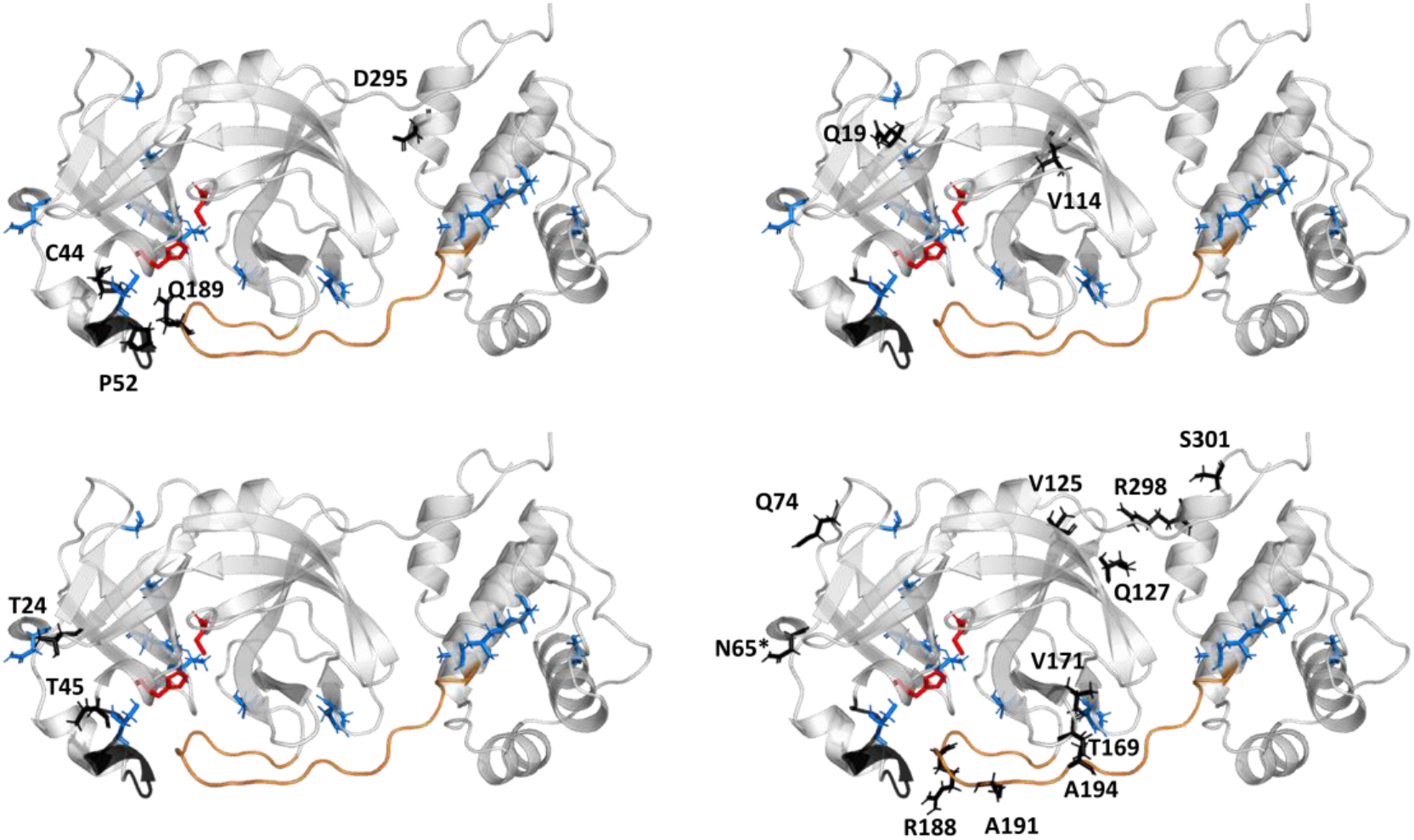
Localisation of the evolutionary-correlated residues of Mpros (black sticks). The CMA analysis provided four groups of evolutionary-correlated residues. The SARS-CoV-2 Mpro structure is presented as cartoon, the active site residues are shown as red sticks, the unique residues of the SARS-CoV-2 Mpro as blue sticks, the asterisk (*) indicates the residue belonging to the evolutionary-correlated residues, unique for SARS-CoV Mpro. The loop C44-P52 is coloured black and the F185-T201 loop is orange. Please note, that within one of the correlated groups (upper left) the residues from C44-P52 loop are correlated with Q189 from the linker loop and with residue from III domain.

In the interest of examining the energetical effect of the 12 amino acid replacement in the SARS-CoV-2 Mpro structure, we calculated their energetic contributions to the protein’s stability using FoldX [34]. As expected, the differences in total energies of the SARS-CoV Mpro and variants with introduced mutation from SARS-CoV-2 Mpro residue did not represent a significant energy change (Supplementary Table S1). The biggest energy reduction was found for mutation H134F (−0.85 kcal/mol) and mutations R88K, S94A, T285A, I286L only slightly reduced the total energy (Supplementary Table S1).

To investigate further possible mutations of SARS-CoV-2 Mpro, single nucleotide substitutions were introduced to the SARS-CoV-2 main protease gene. If a substitution of a single nucleotide caused translation to a different amino acid compared to the corresponding residue in the wild-type structure, an appropriate mutation was proposed with FoldX calculations. The most energetically favourable potential mutations were chosen based on a −1.5 kcal/mol threshold (Figure 7A, Supplementary Table S2). Most of the energetically favourable potential mutations include amino acids that are solvent-exposed on the protein’s surface, according to NetSurfP [35] results. These results show that in general, exposed amino acids are more likely to mutate.

**Figure 7.**
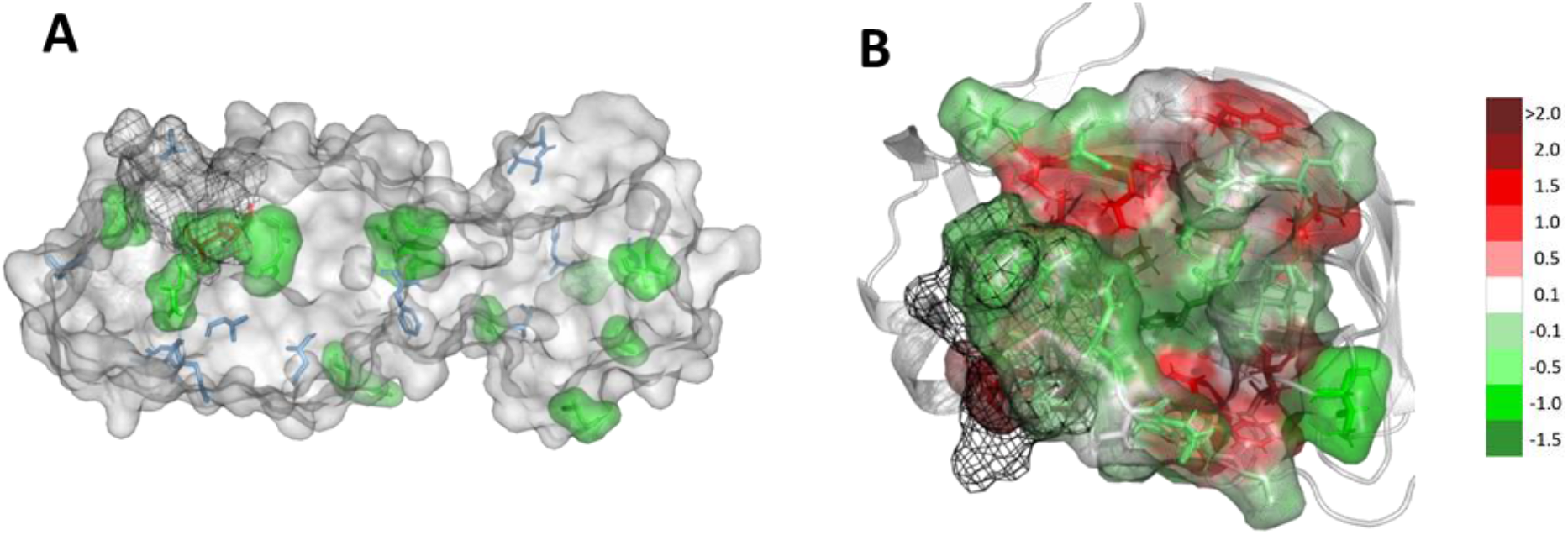
Potential mutability of SARS-CoV-2 Mpro. (A) Structure of SARS-CoV-2 Mpro with the most energetically favourable potential mutations of amino acids marked as green surface. Positions of amino acids that differ from the ones in SARS-CoV Mpro structure marked as blue sticks. Catalytic dyad marked as red. (B) The catalytic site of SARS-CoV-2 Mpro is shown as surface with the most energetically favourable potential mutations shown as green, neutral as white and unfavourable as red. The C44-P52 loop is shown as black mesh.

Additionally, the potential mutability of the binding cavity was investigated. Residues belonging to the binding cavity were found within 7 Å from the N3 inhibitor. Then, we calculated the differences in the Gibbs free energy of protein folding with respect to the wild-type protein (Supplementary Table S3) and presented the results as a heat map. The most energetically favourable potential mutations are shown as green, neutral as white and unfavourable as red (Figure 7B). Interestingly, residues forming the catalytic dyad, namely H41 and C145, are also prone to mutate. However, probably the most important message comes from the analysis of the potential mutability of the C44-P52 loop. Mutation of four of them has a stabilising effect for the protein and for the rest the effect is near-neutral. These results indicate that the future evolution of the Mpro protein can significantly reduce the potential use of this protein as a molecular target for coronavirus treatment due to a highly probable development of drug resistance of this virus through mutations.

## 3. Discussion

As we have shown in previous research, tracking of water molecules in the binding cavity combined with the local distribution approach can identify catalytic water positions [36]. Indeed, despite differences in the size and dynamics of the binding cavities of SARS-CoV and SARS-CoV-2 Mpros, the main identified water was always found in a position next to the H41 residue, and this location is assumed to indicate catalytic water of Mpro replacing the missing third catalytic site amino acid [28]. That was the first quality check of our methodology that approved our approach, and has initiated further investigations.

As reported in the previous research, the overall plasticity of Mpro is required for proper enzyme functioning [37,38]. In the case of SARS-CoV the truncation of the linker loop (F185-T201) gave rise to a significant reduction in protein activity and confirmed that the proper orientation of the linker allows the shift between dimeric and monomeric forms [39]. Dimerisation of the enzyme is necessary for its catalytic activity and the proper conformation of the seven N-terminal residues (N-finger) is required [40]. In SARS-CoV-2 Mpro, the T285 is replaced by alanine, and the I286 by leucine. It has been shown that replacing S284, T285, and I286 by alanine residues in SARS-CoV Mpro leads to a 3.6-fold enhancement of the catalytic activity of the enzyme. This is accompanied by changes in the structural dynamics of the enzyme that transmit the effect of the mutation to the catalytic centre. Indeed, the T285A replacement observed in the SARS-CoV-2 Mpro allows the two domains III to approach each other a little closer [41].

The comparison of MD simulations of both main proteases initiated from different starting conformations (with and without N3 inhibitor) suggests that beside plasticity of the whole protein, there can be large differences between the accessibility to the binding cavity and/or the accommodation of the shape of the cavity in response to the inhibitor that can be bound. There are also differences in the outer pockets’ maximal accessible volumes between the two structures of SARS-CoV main proteases; the apo SARS-CoV Mpro structure used as a starting point of MD simulations has shown the largest MAV of all the analysed systems. These results suggest that the SARS-CoV main proteases’ binding cavity is highly flexible and changes both in volume and shape significantly alter the ligand binding. This finding indicates a serious obstacle for a classical virtual screening approach and drug design in general. Numerous novel compounds that are considered as potential inhibitors of SARS-CoV, have not reached the stage of clinical trials. The lack of success might be related to the above-mentioned plasticity of the binding cavity. Some of these compounds have been used for docking and virtual screening research aimed not only at SARS-CoV [42,43] but also at the novel SARS-CoV-2 [15–21]. Such an approach focuses mostly on the structural similarity between the binding pockets, but ignores the fact that the actual available binding space differs significantly. In general, a rational drug design can be a very successful tool in the identification of possible inhibitors in cases where the atomic resolution structure of the target protein is known. For a new target, when a highly homologous structure is available, a very logical strategy would be seeking chemically similar compounds or creating derivatives of this inhibitor, and finding those compounds that are predicted to have a higher affinity for the new target structure than the original one. This would be expected to work for SARS-CoV-2 proteins (such as Mpro) using SARS-CoV proteins as templates. However, our in-depth analysis indicates a very different situation taking place, with major shape and size differences emerging due to the binding site flexibility. The continuous effort of Diamond Light Source group [44] performing massive XChem crystallographic fragment screen against Mpro supports our finding on large binding site flexibility (Supplementary Figure S4). Therefore, repurposing SARS drugs against COVID-19 may not be successful due to major shape and size differences, and despite docking methods, the enhanced sampling should be considered.

The analysis of the water hot-spots shows that the catalytic water hot-spot dominates water distribution inside the binding cavity. The remaining water hot-spots correspond to a much lower water density level and are on the borders of the binding cavity, which suggests a rather hydrophobic or neutral interior of the binding cavity. The MixMD simulations performed with various cosolvents have further confirmed these observations. The largest number and the densest hot-spots were located within the binding cavity and the region essential for Mpros dimerisation [31], between the II and III domains. The deep insight into the local hot-spots distribution of the various cosolvents underlines the large differences in binding sites plasticity. The smaller binding cavity of the SARS-CoV-2 enlarged significantly in the presence of a highly hydrophobic cosolvent. The benzene hot-spots were detected deep inside the cavity, and also near the C44-P52 loop. In contrast, in the case of the SARS-CoV, benzene hot-spots were located only in the vicinity of the C44-P52 loop. Such a conclusion may also imply that a sufficiently potent inhibitor of SARS-CoV and/or SARS-CoV-2 Mpros needs to be able to open its way to the active site before it can successfully bind to its cavity. These results support the regulatory role of the C44-P52 loop and again alert against unwarranted use of simplified approaches for drug repositioning or docking.

The difficulties in targeting the active site of the Mpros are also explained by evolutionary study and potential mutability analysis. As already pointed out, the C44-P52 loop is likely to regulate the access to the active site by enabling entrance of favourable small molecules and blocking the entry of unfavourable ones. The second important loop, F185-T201, which starts in the vicinity of the binding site and links I and II domains with the III domain contributes significantly to Mpro dimerisation [39].

The initial analysis of the effect of the 12 amino acid replacements in SARS-CoV-Mpro on the SARS-CoV-2 Mpro structure stability was expected to provide neutral or stabilising contribution to proteins folding. Indeed, all replacements were found to stabilise the protein’s folding (e.g., H134F: −0.85 kcal/mol) or have almost neutral character (e.g., R88K, S94A, T285A, I286L). The analysis of the potential risk of further Mpro structure evolution within the binding cavity suggests that mutations of residues that contribute to ligand binding or access to the active site are energetically favourable, and likely to occur. Some of the residues that are prone to mutate would provide the inactive enzyme (e.g., the residues forming the catalytic dyad) and therefore could be considered as a blind alley in enzyme evolution, but others (e.g., amino acids from the C44-P52 loop, T45, S46, E47, L50) could significantly modify the inhibitors binding mode of Mpro. The location of prone to mutate residues on the regulatory loop questions the effort for Mpro active site inhibitors design as a long term strategy. However, our results indicate also residues that are energetically unfavourable to mutate (e.g., P39, R40, P52, G143, G146, or L167), which could provide an anchor for successful drug design that can outlast coronavirus’ Mpros variability in future. Alternatively, we would suggest targeting the region between II and III domains, which contributes to the dimer formation.

## 4. Materials and Methods

### 4.4 Classical MD simulations

The H++ server [45] was used to protonate the SARS-CoV-2 and SARS-CoV main proteases’ structures (PDB IDs: 6lu7, 6y2e, 2amq and 1q2w, respectively) using standard parameters and pH 7.4. The missing 4-amino-acids-long loop of the 1q2w model was added using the corresponding loop of the 6lu7 model. Water molecules were placed using the combination of 3D-RISM [46] and the Placevent algorithm [47]. The AMBER 18 LEaP [48] was used to immerse models in a truncated octahedral box of TIP3P water molecules and prepare the systems for simulation using the ff14SB force field41. Additionally, 4 and 3 Na + ions were added to the SARS-CoV-2 and the SARS-CoV, respectively. AMBER 18 software [48] was used to run 2 μs simulations of both systems. The minimisation procedure consisted of 2000 steps, involving 1000 steepest descent steps followed by 1000 steps of conjugate gradient energy minimisation, with decreasing constraints on the protein backbone (500, 125 and 25 kcal x mol-1 x Å2) and a final minimisation with no constraints of conjugate gradient energy minimization. Next, gradual heating was performed from 0 K to 300 K over 20 ps using a Langevin thermostat with a temperature coupling constant of 1.0 ps in a constant volume periodic box. Equilibration and production stages were run using the constant pressure periodic boundary conditions for 1 ns with 1 fs step and 50 ns with a 2 fs time step, respectively. Constant temperature was maintained using the weak-coupling algorithm for 50 ns of the production simulation time, with a temperature coupling constant of 1.0 ps. Long-range electrostatic interactions were modelled using the Particle Mesh Ewald method with a non-bonded cut-off of 10 Å and the SHAKE algorithm. The coordinates were saved at an interval of 1 ps.

### 4.2. Mixed-solvent MD simulations – cosolvent preparation

Six different cosolvents: acetonitrile (ACN), benzene (BNZ), dimethylsulfoxide (DMSO), methanol (MEO), phenol (PHN), and urea (URE) were selected to perform the mixed-solvent MD simulations. The chemical structures of cosolvents molecules were downloaded from the ChemSpider database [49] and a dedicated set of parameters was prepared. Parameters for CAN were adopted from the work by Nikitin and Lyubartsev [50], and parameters for URE were modified using the 8Mureabox force field to obtain parameters for a single molecule. For the rest of the co-solvent molecules, parameters were prepared using Antechamber [51] with Gasteiger charges [52].

### 4.3. Mixed-solvent MD simulations – initial configuration

The Packmol software [53] was used to build the initial systems consisting of protein (protonated according to the previously described procedure), water, and particular cosolvent molecules. 4 and 3 Na^+^ ions were added to the SARS-CoV-2 Mpro and the SARS-CoV Mpros, respectively. It was assumed that the percentage concentration of the cosolvent should not exceed 5% (in the case of ACN, DMSO, MEO, and URE), or should be about 1% in the case of BNZ and PHN phenol (see Supplementary Table S4). The mixed-solvent MD simulation procedures (minimization, equilibration, and production) carried out using the AMBER 18 package were identical as for the classical MD simulations. Only the heating stage differed – it was extended up to 40 ps.

### 4.4. Water and cosolvent molecules tracking

The AQUA-DUCT 1.0 software was used to track water and cosolvent molecules. Molecules of interests, which have entered the so-called *Object,* defined as 5Å sphere around the centre of geometry of active site residues, namely H41, C145, H164, and D187, were traced within the *Scope* region, defined as the interior of a convex hull of both COVID-19 Mpro and SARS Mpro Cα atoms. All visualisations were made in PyMol [54].

AQUA-DUCT was used to analyse maximal accessible volume (MAV), defined as the *outer* pockets; the *outer* pocket represents the maximal possible space that could be explored by tracked molecules.

### 4.5. Hot-spot identification and selection

AQUA-DUCT was used to detect regions occupied by molecules of interests, and identify the densest sites using a local solvent distribution approach. Those so-called hot-spots could be calculated as local and/or global, based on the distribution of tracked molecules which visited the *Object* (local) or just the *Scope* without visiting the *Object* (global); here, they are considered as potential binding sites. For clarity, the size of each sphere representing a particular hot-spot has been changed to reflect its occupation level. The selection of the most significant hot-spots consisted of indicating points showing the highest density in particular regions. From the set of points in the space, small groups of hot-spots were determined. Groups were further defined by distance (radius) from each other. Any point found within a distance shorter than the determined radius (3Å) from any other point being part of a given group was counted toward the group. For each so designated group of points, one showing the highest density was chosen as representing the place.

### 4.6. Obtaining SARS-CoV-2 Mpro gene sequence

SARS-CoV-2 Mpro was downloaded from the PDB as a complex with an N3 inhibitor (PDB ID: 6lu7). Tblastn [55] was run based on the protein amino acid sequence. 100% identity with 10055-10972 region of SARS-CoV-2 Mpro complete genome (Sequence ID: MN985262.1) was obtained. Blastx [56] calculations were run with the selected region, and orf1a polyprotein (NCBI Reference Sequence: YP_009725295.1) amino acid sequence, identical with the previously downloaded SARS-CoV-2 Mpro, was received.

### 4.7. Energetic effect of amino acids substitutions

FoldX software was used to insert substitutions into the structures of SARS-CoV and SARS-CoV-2 Mpros. To analyse the changes in the two structures, 12 single-point mutations were introduced to the SARS structure. Each of the residues in SARS-CoV Mpro was mutated to the respective SARS-CoV-2 Mpro residue, and the difference in total energies of the wild-type SARS-CoV-2 Mpro and the mutant structures were calculated. Then, to investigate further possible mutations of SARS-CoV-2 Mpro, single nucleotide substitutions were introduced to the SARS-CoV-2 main protease gene. If a substitution of a single nucleotide caused translation to a different amino acid than the corresponding residue in the wild-type structure, an appropriate mutation was proposed with FoldX software.

### 4.8 Comulator calculations of correlation between amino acids

SARS-CoV Mpro was downloaded from the PDB (PDB ID: 1q2w). Blast [57] was run based on the amino acid sequence. As a result, 2643 sequences of viral main proteases similar to chain A SARS-CoV Mpro were obtained. Clustal Omega [58] was used to prepare an alignment of those sequences. Comulator was then employed to calculate the correlation between amino acids and based on the results, groups of positions in SARS-CoV Mpro sequence were selected whose amino acid occurrences strongly depended on each other.

## 5. Conclusions

In this paper, we reported on molecular dynamics simulations of the main protease (Mpro), whose crystal structures have been released. We compared the Mpro for SARS-CoV-2 with a highly similar SARS-CoV protein. In spite of a high level of sequence similarity between these two homologous proteins, their active sites show major differences in both shape and size indicating that repurposing SARS drugs for COVID-19 may be futile. Furthermore, a detailed analysis of the binding pocket’s conformational changes during simulation time indicates its flexibility and plasticity, which dashes hope for rapid and reliable drug design. Moreover, our findings show the presence of a flexible loop regulating the access to the binding site pocket. A successful inhibitor may need to have an ability to relocate the loop from the entrance to bind to the catalytic pocket. However, mutations leading to changes in the amino acid sequence of the loop, while not affecting the folding of the protein, may result in the putative inhibitors’ inability to access the binding pocket and provide a probable development of drug resistance. To avoid that situation that the future evolution of the Mpros can wipe out all our efforts, we should focus on key functional residues or those whose further mutation will destabilise the protein.

## Supplementary Materials

Supplementary materials are available

## Author Contributions

Conceptualization, A.G.; methodology, M.B., K.M. A.R and A.S.; software, M.B., K.M. A.R and A.S.; validation, M.B., K.M. and A.G.; formal analysis, M.B., K.M. A.R and A.S.; investigation, M.B., K.M. A.R and A.S.; resources, M.B and K.M.; data curation, M.B and K.M.; writing—original draft preparation, M.B and K.M.; writing—review and editing, J.A.T. and A.G.; visualization, M.B., K.M. A.R and A.G; supervision, A.G.; project administration, A.G.; funding acquisition, J.A.T. and A.G. All authors have read and agreed to the published version of the manuscript.

## Abbreviations

Mpro: Main protease
CoVs: Coronaviruses
ORFs: Open reading frames
3CLpro: Chymotrypsin-like cysteine protease
S: Spike surface glycoprotein
E: Small envelope protein
M: Matrix protein
N: Nucleocapsid protein
N3: N-[(5 methylisoxazol-3-yl)carbonyl] alanyl-L-valyl-N~1-((1R,2Z)-4-(benzyloxy)-4-oxo-1--{[(3R)-2-oxopyrrolidin-3-yl]methyl}but-2-enyl)-L-leucinamide
cMD: Classical molecular dynamics simulations
MixMD: Mixed-solvent molecular dynamics simulations
PDB: Protein Data Bank
MAV: Maximal accessible volume
ACN: Acetonitrile
BNZ: Benzene
DMSO: Dimethylsulfoxide
MEO: Methanol
PHN: Phenol
URE: Urea
CMA: Correlated mutation analysis

