## Supplementary Material for "Structural and Evolutionary Analysis Indicate that the SARS-CoV-2 Mpro is an Inconvenient Target for Small-Molecule Inhibitors Design"

Supplementary Table S1. Differences between SARS-CoV-2 and SARS-CoV Mpros proteins. The last column shows differences in total energies (in kcal/mol) calculated as differences in Gibbs free energy folding between SARS-CoV Mpro and introduced single-point mutations as they are in SARS-CoV-2 Mpro structure.

| ID | SARS-CoV-2 Mpro | SARS-CoV Mpro | domain | buried/exposed (based on the NetSurfP calculations) | total energy differences [kcal/mol] (based on the FoldX calculations) |
|---|---|---|---|---|---|
| 35 | V | T | I | B | 0.90 |
| 46 | S | A | I | B | 0.14 |
| 65 | N | S | I | B(SARS-CoV-2)/ E(SARS-CoV) | 0.32 |
| 86 | V | L | I | B | 2.76 |
| 88 | K | R | I | E | -0.42 |
| 94 | A | S | I | E | -0.41 |
| 134 | F | H | II | E | -0.85 |
| 180 | N | K | II | E | 1.29 |
| 202 | V | L | III | B | 1.78 |
| 267 | S | A | III | B | 2.18 |
| 285 | A | T | III | E | -0.37 |
| 286 | L | I | III | E | -0.08 |
| Changes in the protein's sequence, not present in the crystallographic structure | | | | | |
| 305 | F | Q | | E | |
| 306 | Q | G | | E | |

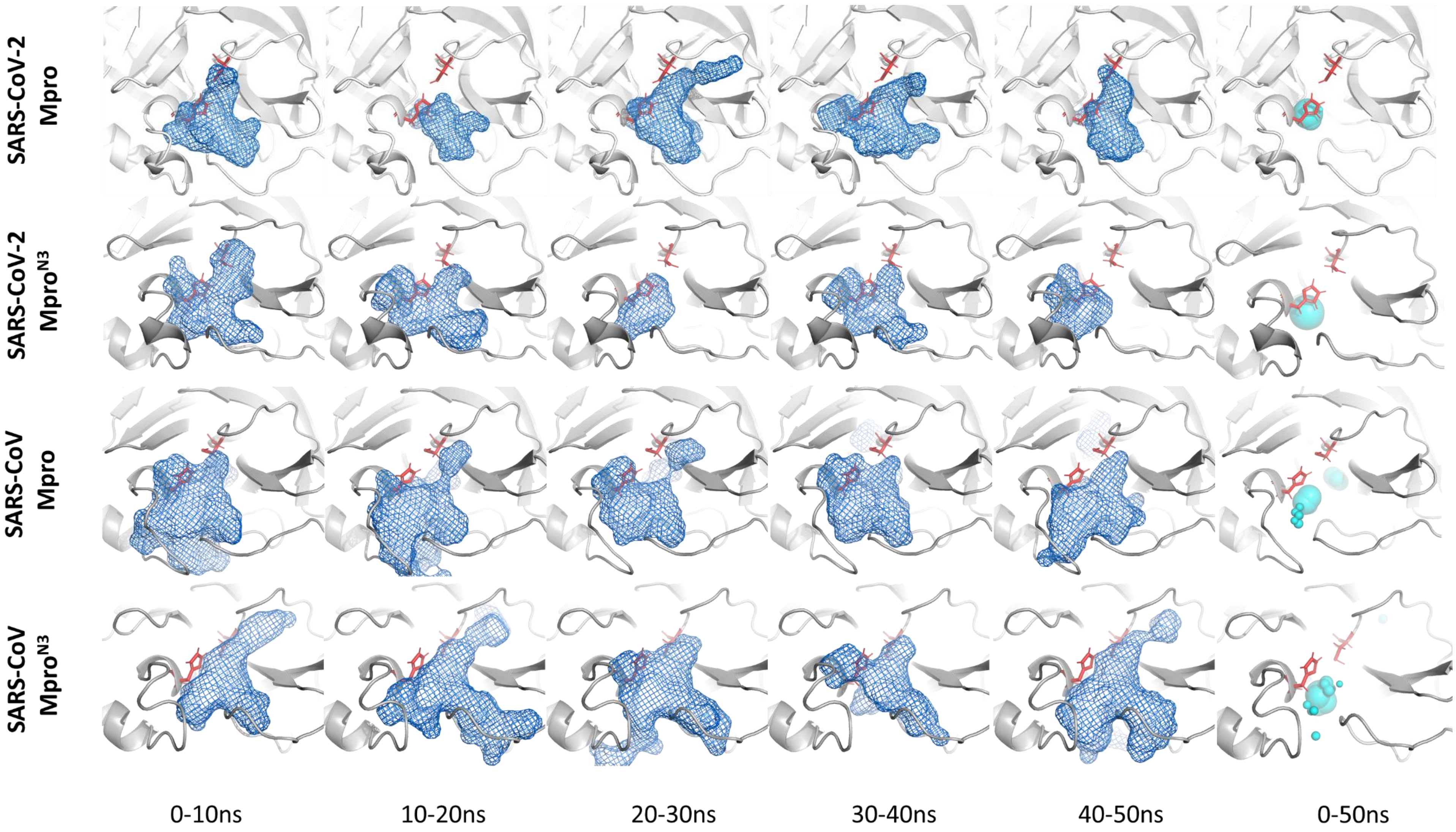


Supplementary Figure1. The example of time mode analysis of the maximal accessible volume (MAV) (blue mesh) of Mpro structures. The catalytic dyad is shown as red sticks. The last column shows the average location of water hot-spots (cyan spheres) during the simulation time. The position of the biggest hot-spot in each row reflects the position of the catalytic water molecule.

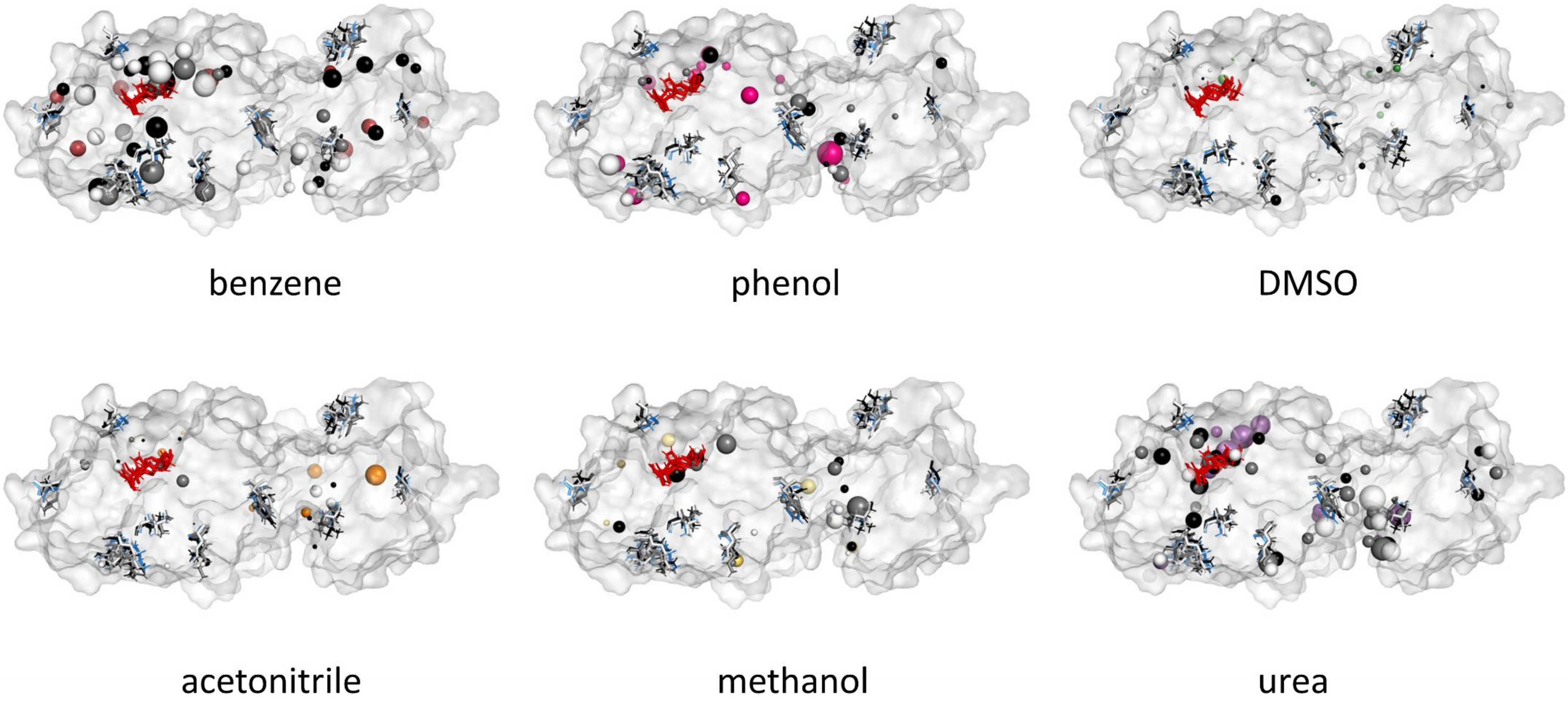


Supplementary Figure S2. Localisation of the global hot-spots of all analysed Mpros. SARS-CoV Mpro$^{N3}$ and SARS-CoV Mpro. The structures of all analysed Mpro structures are superposed and the colour-coding is as follows: orange, red, yellow, green, pink and purple hot-spots are from the SARS-CoV-2 Mpro$^{N3}$, white hot-spots from the SARS-CoV-2 Mpro, black hot-spots from the SARS-CoV Mpro$^{N3}$, and grey hot-spots from the SARS-CoV Mpro structure. The active site residues are shown as red sticks, the differing residues of the SARS-CoV-2 Mpro as blue sticks, and the proteins' structures are shown in surface representation.

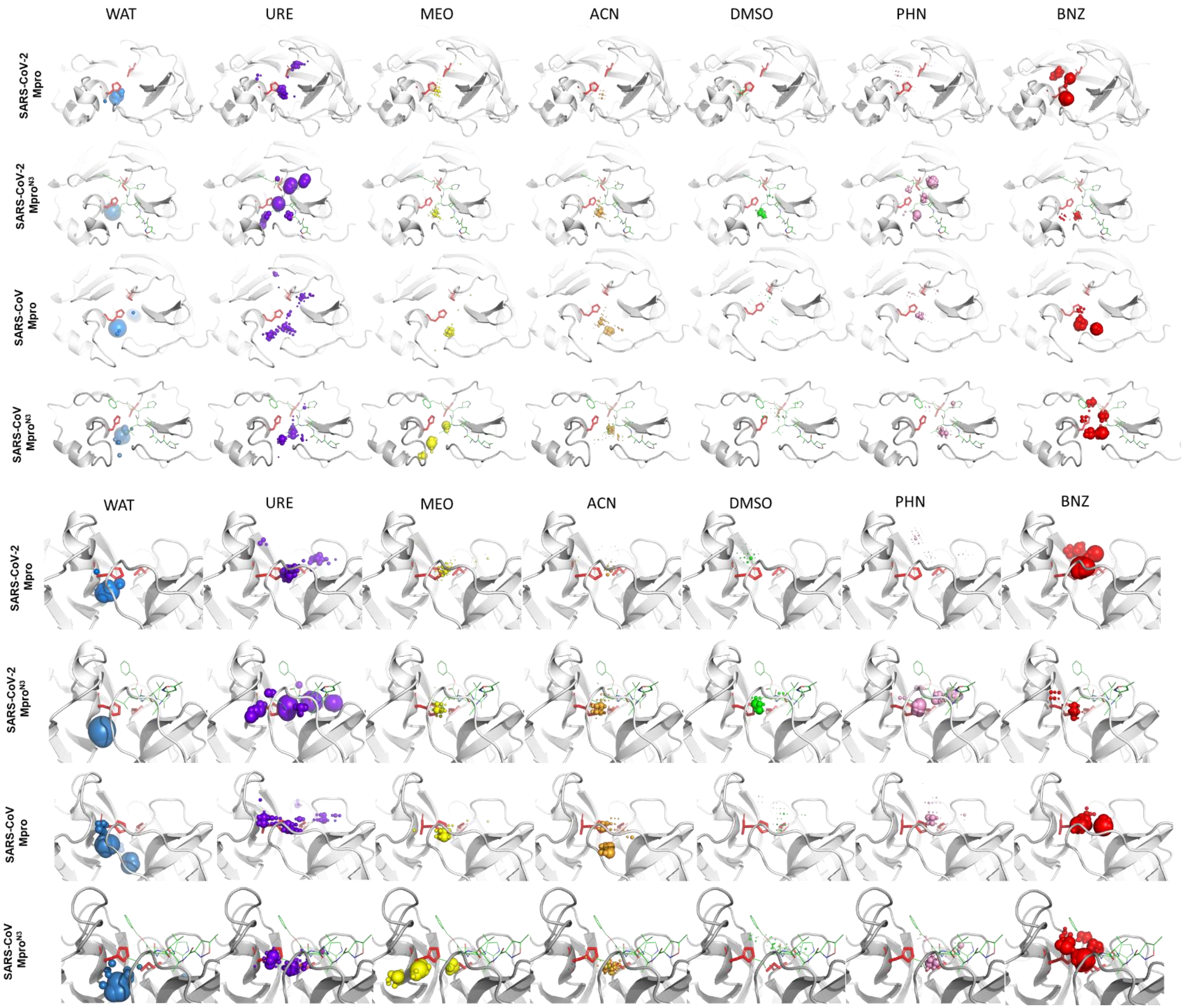


Supplementary Figure S3. Localisation of the local hot-spots of all analysed Mpros: COVID-19 Mpro, SARS-CoV Mpro and SARS-CoV Mpro-f. Hot-spots for individual cosolvents are represented by spheres, and their size reflects the hot-spots density. The colour-coding is as follows: purple - urea, green - DMSO, yellow - methanol, orange - acetonitrile, pink - phenol, red - benzene. The active site residues are shown as red sticks, the N3 inhibitor structure as green lines, and the proteins' structures are shown in cartoons representation.

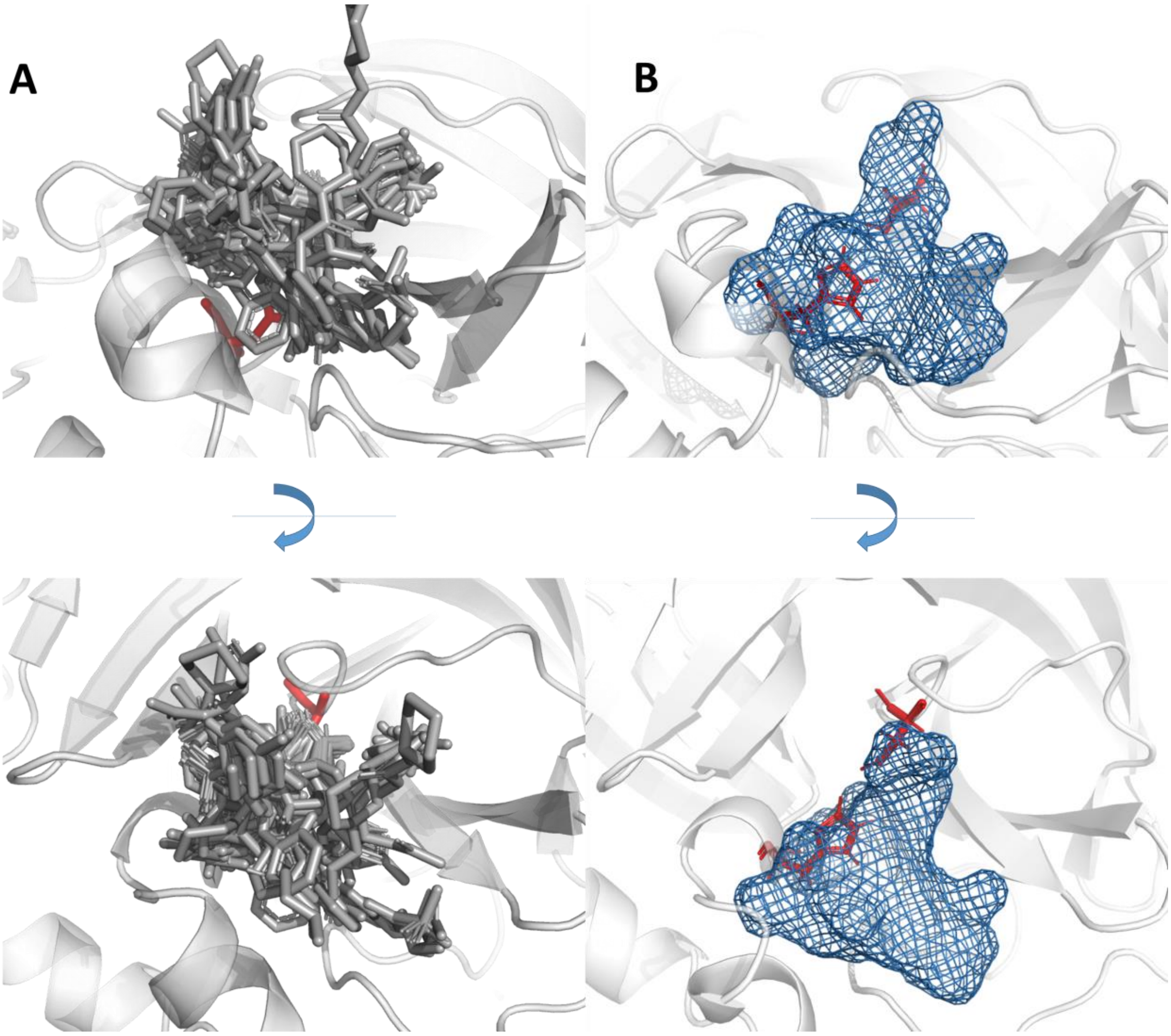


Supplementary Figure S4. Comparison of the space occupied by covalently bound fragments in the active site cavity (Diamond Light Source group) (A) with maximal accessible volume calculated by AQUA-DUCT software (B). Please note, that part of the ligands extends beyond protein surface.

Supplementary Table S2. FoldX results for the most energetically favourable potential mutations in the SARS-CoV-2 Mpro structure. Amino acids from the binding cavity are marked **bold**.

| Mutation | Energy difference [kcal/mol] | buried/exposed* |
|---|---|---|
| A260D | -3.67 | E |
| Y154H | -2.04 | E |
| T21I | -2.01 | B |
| **H41L** | **-1.95** | **B** |
| Q127L | -1.89 | E |
| A194P | -1.82 | E |
| A129V | -1.75 | B |
| Q306R | -1.73 | E |
| **H164L** | **-1.71** | **B** |
| S301L | -1.68 | E |
| V233L | -1.58 | B |
| Q244P | -1.54 | E |
| N53D | -1.51 | E |

* based on the NetSurfP calculations

Supplementary Table S3. FoldX results for binding cavity amino acids (7Å within the N3 inhibitor). Catalytic dyad is marked **bold**.

| Mutation | Energy [kcal/mol] | buried/exposed* |
|---|---|---|
| **H41L** | **-1.95** | **B** |
| H164L | -1.71 | B |
| T169I | -1.49 | E |
| T45S | -0.93 | E |
| E47Q | -0.91 | E |
| T24R | -0.84 | E |
| S46P | -0.81 | B |
| Q189L | -0.79 | E |
| T26K | -0.78 | E |
| **C145F** | **-0.77** | **B** |
| A191P | -0.58 | E |
| H163Y | -0.55 | B |
| E166Q | -0.43 | E |
| L50R | -0.37 | E |
| A193T | -0.36 | E |
| H172L | -0.35 | B |
| D187A | -0.35 | E |
| V186F | -0.30 | E |
| L141R | -0.28 | B |
| N142Y | -0.22 | B |
| N119Y | -0.08 | E |
| S144A | -0.08 | B |
| A173V | -0.02 | B |
| T190I | 0.05 | E |
| Q192L | 0.22 | E |
| S147T | 0.29 | B |
| N28I | 0.37 | B |
| Y54F | 0.40 | B |
| C44S | 0.41 | B |
| M49L | 0.42 | B |
| V42L | 0.74 | B |
| Y118F | 0.78 | B |
| T25A | 0.84 | E |
| L27I | 1.04 | B |
| F140Y | 1.09 | B |
| F181Y | 1.37 | B |
| F185L | 1.47 | E |
| P52A | 1.54 | E |
| L167I | 2.30 | B |
| G143R | 2.42 | B |
| P39A | 3.21 | B |
| R40I | 3.50 | E |
| G146A | 9.72 | B |

* based on the NetSurfP calculations

Supplementary Table S4. The final percentage concentration of particular cosolvents for both SARS-CoV-2 and SARS-CoV Mpros systems.

| Cosolvent | Concentration [%] | Number of added molecules | | | |
|---|---|---|---|---|---|
| | | SARS-CoV-2 Mpro$^{N3}$ | SARS-CoV-2 Mpro | SARS-CoV Mpro$^{N3}$ | SARS-CoV Mpro |
| ACN | 4.5 | ACN: 450<br>WAT: 19712 | ACN: 450<br>WAT: 19924 | ACN: 450<br>WAT: 19801 | ACN: 450<br>WAT: 19858 |
| BNZ | 1.0 | BNZ: 50<br>WAT: 19712 | BNZ: 50<br>WAT: 19924 | BNZ: 50<br>WAT: 19801 | BNZ: 50<br>WAT :19858 |
| DMSO | 4.8 | DMSO: 250<br>WAT: 19712 | DMSO: 250<br>WAT: 19924 | DMSO: 250<br>WAT: 19801 | DMSO: 250<br>WAT: 19858 |
| MEO | 4.3 | MEO: 550<br>WAT: 19712 | MEO: 550<br>WAT: 19924 | MEO: 550<br>WAT: 19801 | MEO: 550<br>WAT: 19858 |
| PHN | 1.2 | PHN: 50<br>WAT: 19712 | PHN: 50<br>WAT: 19924 | PHN: 50<br>WAT: 19801 | PHN: 50<br>WAT: 19858 |
| URE | 4.4 | URE: 300<br>WAT: 19712 | URE: 300<br>WAT: 19924 | URE: 300<br>WAT: 19801 | URE: 300<br>WAT: 19858 |
